# Tumor cell-derived lymphotoxin alpha triggers metastatic extravasation through TNFRs/cIAP1

**DOI:** 10.1101/766485

**Authors:** Lazaros Vasilikos, Kay Hänggi, Lisanne M. Spilgies, W. Wei-Lynn Wong

**Affiliations:** Institute of Experimental Immunology, University of Zurich, CH-8057 Zurich, Switzerland

**Keywords:** cIAPs, Smac mimetics, permeability, tumor extravasation, endothelial cell, lymphotoxin A, TNF receptors

## Abstract

Metastasis involves the interaction of the tumor, immune and endothelial cells. Cell death proteins, such as inhibitors of apoptosis proteins (IAPs), are critical players in survival, inflammation and permeability. Whether the use of Smac mimetics, which target cIAP1/2 for degradation would affect metastasis is unknown. We show Smac mimetics reduced metastasis due to the loss of cIAP1 but not cIAP2 in experimental metastasis models. The endothelial compartment rather than the immune cells was responsible for reduction of extravasation upon loss of cIAP1. Loss of cIAP1 in primary endothelial cells did not lead to cell death but resulted in an unresponsive endothelium barrier to permeability factors causing a reduction in tumor cell extravasation. Unexpectedly, the co-loss of TNFR1 and cIAP1 restored the tumor load. We were surprised to find lymphotoxin alpha (LTA), and not TNF, secreted by the tumor cells was critical for the extravasation. Using TCGA data, we found high levels of LTA mRNA expression correlated with decreased survival in kidney carcinoma and associated with advance disease stage. Our data suggest that Smac mimetics, targeting cIAP1/2, may reduce metastasis to the lung through a LTA/TNFR mechanism by altering the endothelial barrier and inhibiting the ability of tumor cells to extravasate.

## Introduction

Elevated levels of cIAPs (cIAP1/2), in part due to genomic amplification, have been reported in a variety of human cancers such as hepatocellular carcinoma(1), lung cancer(2) and esophageal squamous cell carcinoma(3). In cervical squamous cell carcinoma, elevated levels of cIAP1 are correlated with chemoresistance to radiotherapy(4) and in colorectal and bladder cancer elevated levels of cIAPs are correlated with advanced stages of tumors and poor survival(3, 4).

Because of the role of IAPs to inhibit cell death, pharmaceutical companies developed synthetic compounds to mimic the naturally inhibitor of IAPs, Smac/DIABLO(5, 6). The release of Smac from the mitochondria results in autoubiquitination and degradation of cIAP1 and cIAP2 by binding to their BIR3 domain(5, 6) and inhibition of XIAP and caspases through the binding of the BIR2 and BIR3 domain of XIAP(7). Likewise, Smac mimetics cause the proteasomal degradation of cIAP1/2 and activation of the alternative NF-κB pathway. NF-κB drives TNF production and in the absence of cIAP1, TNFR1 activation leads to the formation of complex II and subsequent cell death (8–11). TNFR2 activation in the absence of IAPs also induces cell death through a TNF/TNFR1 mechanism(12, 13).

Numerous studies have implicated Smac mimetics in modifying the immune system to cause tumor cell death and reduce tumor burden alone or in combination with known immune modulators(14–18). Moreover, in a B16-F10 subcutaneous tumor model, delivery of Smac mimetic in the tumor site disrupted neo-angiogenesis due to cell death which was rescued by the co-loss of TNFR1/2(19). These results suggest that IAP inhibitors may be used to target the primary tumor, although past clinical trials with IAP inhibitors as single agents have shown poor efficacy.

The majority of patients however, die not due to the primary tumor but because of metastasis. For tumor cells to metastasize, the cells must enter the blood stream, survive within the circulatory system and then extravasate at a secondary site(20). Developmental studies show that the loss of cIAP1/2 results in hemorrhage and loss of endothelial cell integrity(21, 22). Tumor cell-induced endothelial cell death has been implicated as a mode of metastasis(23). Alternatively, loss of cell death proteins in endothelial cells reduced permeability but not viability suggesting a non-cell death function for cell death proteins in endothelial cells(24, 25). Whether Smac mimetics would compromise endothelial cell integrity past development and enhance metastasis is unknown.

Here, we show that the loss of cIAP1 but not cIAP2 in the endothelium obstructs tumor cells extravasation into the lung. Surprisingly, there is no evidence of cell death in this process and the results can be re-capitulated using Smac mimetics (Birinapant). Combined loss of TNFR1 and cIAP1 reverses this phenotype, allowing tumor cells to extravasate into the lung. Our data show that LTA secreted from the tumor cell requires cIAP1 expression in the endothelium to induce tumor cell extravasation. These results suggest that application of Smac mimetics in the clinics might not only contribute to tumor killing by activating the immune system but also prevent tumor metastasis of individual cancer types.

## Results and Discussion

### Smac mimetics decreased tumor load in the lung through reduction of cIAP1 but not cIAP2

To determine if Smac mimetics would alter metastasis, we chose a model that allowed us to focus on the latter part of metastasis. Tumor cells injected intravenously, circulate to the lung, extravasate and form tumor nodules. Birinapant, a Smac mimetic in Phase II clinical trials, was used to target cIAP1/2. Injection of Birinapant led to cIAP1 loss within 1 hour as shown in total lung homogenates and remained reduced up to 48 hours (Figure 1A). Interestingly, treatment with Birinapant (72 and 24 hours prior to injection of B16-F10 melanoma cells) resulted in a significant reduction in lung tumor load at day 13 post tumor challenge (Figure 1B). Injection of Birinapant 6 hours prior to injection of tumor cells, however, did not affect tumor load in the lung (Supplemental Figure 1A). Although Smac mimetics have been shown to cause tumor cell death *in vivo* because of exogenous TNF production from innate immune cells(14, 16), B16-F10 cells were resistant to TNF-induced cell death even in the presence of Smac mimetics (Supplemental Figure 1B).

**Figure 1.**
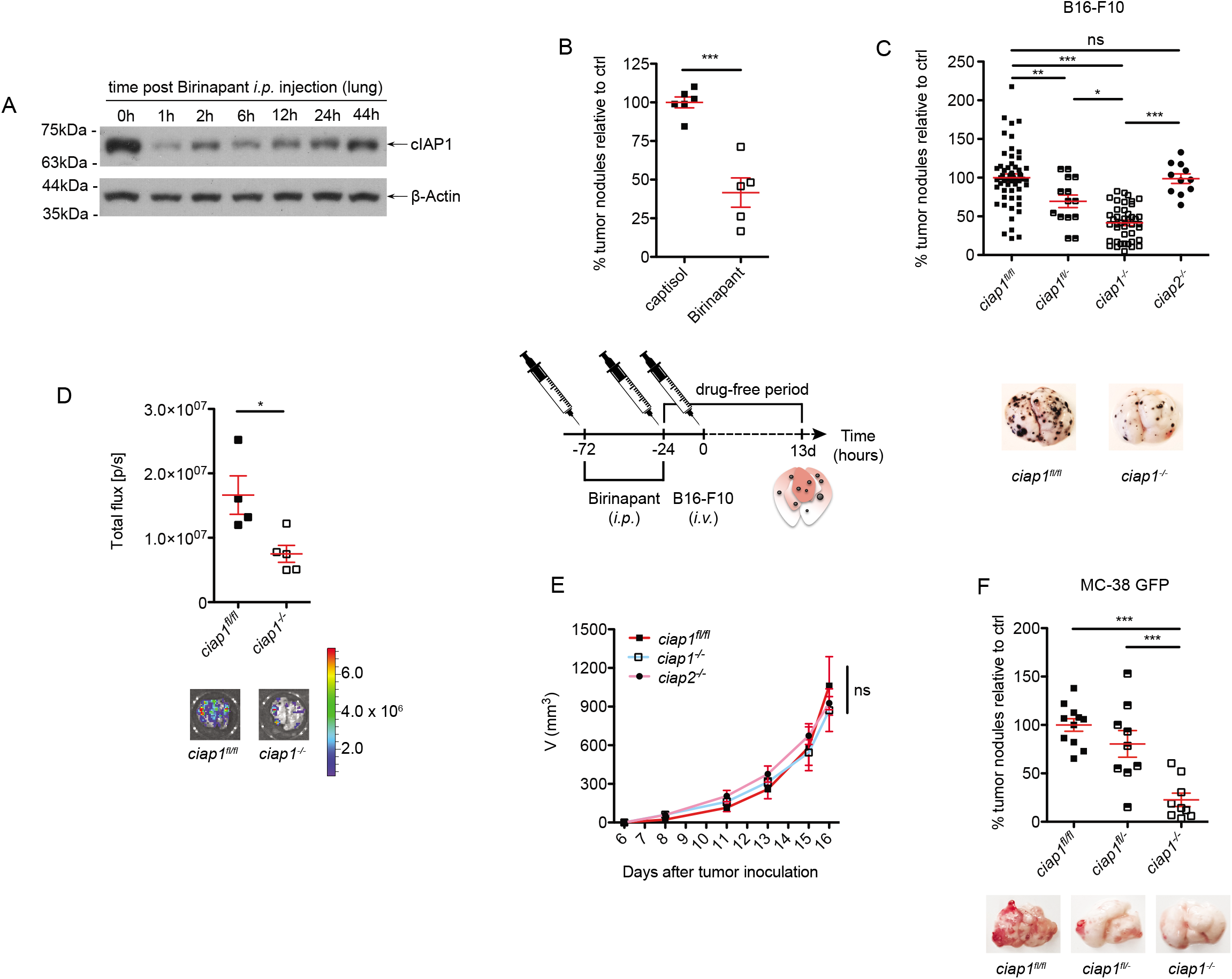
Birinapant impedes metastasis into the lung by causing cIAP1 degradation. A) Total lung lysates from wt mice injected with Birinapant (*i.p.*) for various time points were analysed by western blotting to evaluate the levels of cIAP1. Representative immunoblot of n = 2 experiments. B) Percentage of superficial B16-F10 tumor nodules normalized to the average of the captisol control. Wildtype mice were injected i.p. with Birinapant (Bir, 5mg/kg) or captisol before the tumor challenge. Tumor cells were injected *i.v*. and nodules formed in the lung were counted 13 days later (n = 2 experiments). The experimental set up for Smac mimetic delivery is depicted below the graph. C) Percentage of superficial B16-F10 tumor nodules normalized to the average of the wt control. Tumor cells were injected *i.v*. and nodules formed in the lung were counted 13 days later (n ≥3 experiments). Representative pictures of tumor-bearing lungs are shown. D) B16-F10 cells expressing luciferase were injected *i.v*. and bioluminescence from the lungs was measured 13 days later utilizing the IVIS technology. Representative bioluminescence pictures are shown. E) Subcutaneous tumor growth of B16-F10 cells. Mice were injected subcutaneously with 100,000 tumor cells in the right flank and tumor growth was assessed using calipers (n = 2 experiments, 3-8 mice/group). F) Percentage of superficial MC-38 GFP tumor nodules normalized to the average of the wt control. Tumor cells were injected *i.v*. and nodules formed in the lung were counted 21 days later (n = 2 experiments). Representative pictures of tumor-bearing lungs are shown. Data are presented as mean ± SEM. *p < 0.05; **p < 0.01; ***p < 0.001, ns = not significant. Two-tailed unpaired t-test (B, D), 1-way ANOVA with Bonferroni’s test (C, F), 2-way ANOVA with Bonferroni’s test (E).

To identify whether the loss of cIAP1 or cIAP2 played a dominant role in reducing the number of tumor nodules, we utilized *ciap1^−/−^ciap2^frt/frt^* (*ciap1*^−/−^) and *ciap1^fl/fl^ciap2^−/−^* (*ciap2^−/−^*) mice to mimic the effect of Smac mimetics on the host alone. Approximately 50% fewer tumor nodules were identified in the lungs of *ciap1^−/−^* compared to *ciap1^fl/fl^ciap2^frt/frt^* (*ciap1^fl/fl^*) mice, whereas *ciap1^fl/−^* mice showed an intermediate phenotype (Figure 1C). However, the tumor load in *ciap2^−/−^* mice was similar to widltype mice, suggesting the loss of cIAP1 was the reason for the reduction in tumor nodule numbers. To assess the tumor load in an unbiased manner, B16-F10 luciferase cells were utilized and a significant reduction in bioluminescence intensity was observed in *ciap1^−/−^* mice, correlating with the tumor nodule counts (Figure 1D). Because a previous study using the B16-F10 tumor model showed reduction in subcutaneous tumor growth upon Smac mimetic treatment (Compound A, targeting cIAP1/2 and XIAP)(19) and to ensure tumor cells were not immunologically rejected in *ciap1^−/−^* mice, we injected B16-F10 tumor cells subcutaneously. No difference in tumor growth kinetics among *ciap1^fl/fl^, ciap1^−/−^* or *ciap2^−/−^* mice was observed (Figure 1E). To determine if tumor nodules formed in the lung by other syngeneic tumor lines may be affected by cIAP1 loss in the tumor microenvironment, MC-38 GFP colon carcinoma cells or Lewis lung carcinoma cells (LLC) were injected intravenously and 21 days post injection, the number of tumor nodules were counted. We observed a 75% reduction in MC-38 GFP tumor nodules in the lungs of *ciap1^−/−^* compared to wildtype mice, while no reduction in tumor nodules was seen when LLC cells were injected (Figure 1F and Supplemental Figure 1C). These data suggest the ability of Smac mimetics to reduce the metastatic potential of individual cancer types to the lung by targeting cIAP1 for degradation in the host.

### Loss of cIAP1 in the radio-resistant but not the hematopoietic compartment causes lung tumor load decrease

The extravasation process during metastasis is aided by immune and stroma cells(26, 27). Therefore, we assessed innate immune cell infiltrates in the lung at different time points post tumor cell injection. Two important populations aiding tumor cell extravasation, the inflammatory monocytes/macrophages (Ly6C^+^) and neutrophils (Ly6G^+^), were similarly recruited to the lung at 2h upon tumor challenge in both wildtype and *ciap1^−/−^* mice. Other populations of the innate and adaptive immunity were assessed showing no major changes in kinetics, except from CD8^+^ T cells at the naïve stage. (Figure 2A, Supplemental Figure 2A). Likewise, changes in extracellular remodeling factors, consisting of matrix metalloproteinase (MMP)-related factors, adhesion molecules and cytokines/chemokines were comparable between *ciap1^−/−^* and wildtype mice. Interestingly, MMP-12 (macrophage metalloelastase) has been shown to suppress lung metastases growth(28). In agreement, MMP-12 was detected in high amounts in the lung of *ciap1^−/−^* mice even in the unchallenged state (Figure 2B and Supplemental Figure 2B-2D). Taken together, the recruitment of immune cells at the extravasation site and re-modelling of the extracellular matrix is not compromised upon loss of cIAP1, suggesting the immune system is a redundant factor influencing tumor metastasis in the absence of cIAP1.

**Figure 2.**
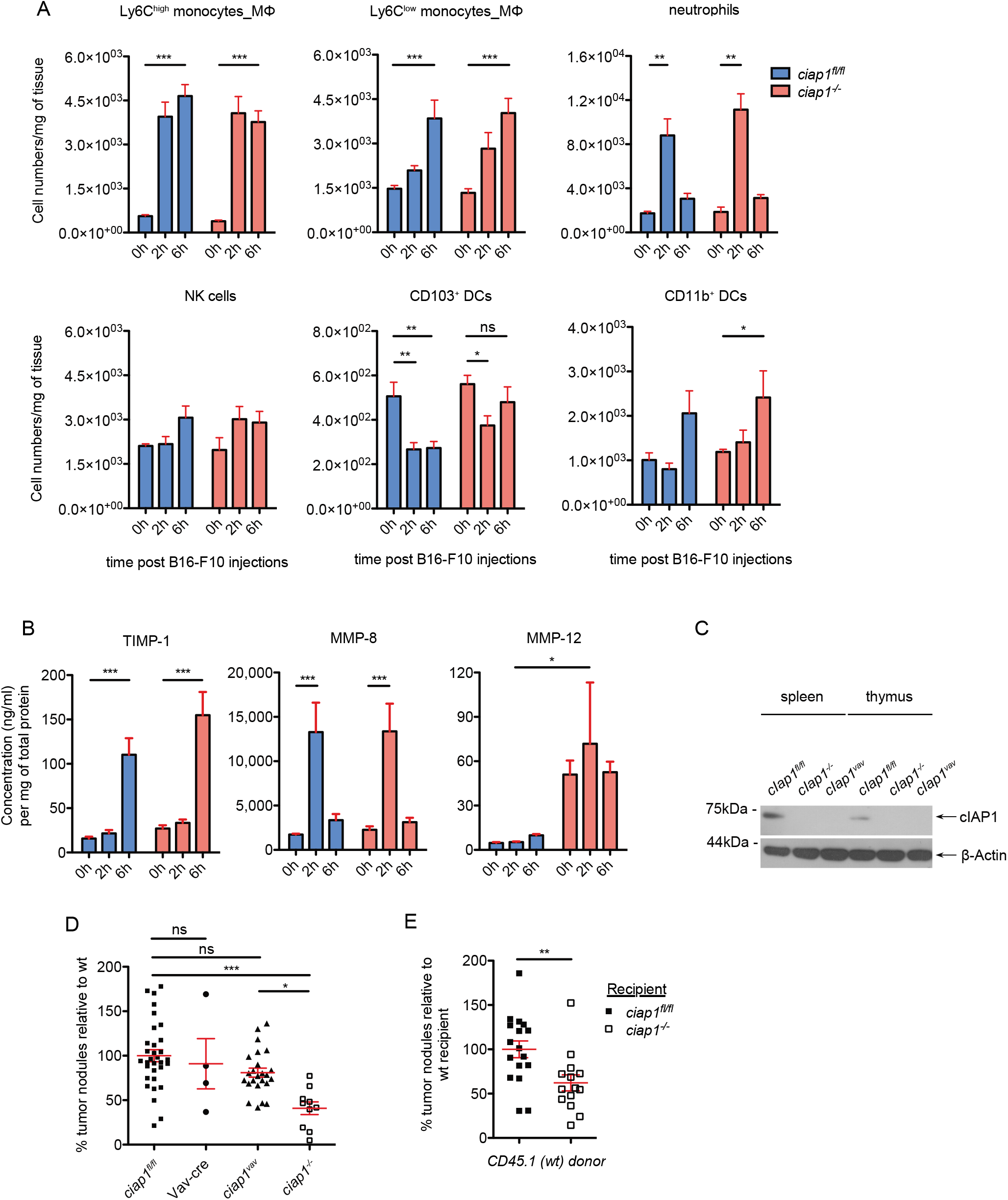
Loss of cIAP1 in the hematopoietic compartment does not alter the lung tumor load. A) Analysis of lung immune cell infiltrates following B16-F10 tumor challenge by flow cytometry. Inflammatory (CD11b^high^MHCII^−^SiglecF^−^Ly6G^−^Ly6C^high^), patrolling monocytes (CD11b^high^MHCII^−^SiglecF^−^Ly6G^−^Ly6C^low^), neutrophils (CD11b^+^Ly6G^+^), natural killer (NK) cells (CD3^−^NK1.1^+^), CD103^+^ DCs (CD11b^int^CD11c^+^CD24^+^CD103^+^) and CD11b^+^ DCs (CD11b^high^MHCII^+^CD24^+^) were pre-gated on singlets, live and CD45+ cells (n = 2 experiments, 6 mice/group). B) Levels of matrix metalloproteinase-related factors assessed using multiplex analysis assay of total lung lysates (n = 2 experiments, 6 mice/group). C) Spleen and thymus, composed mainly by blood cells, from ciap1vav mice were analysed by western blotting to evaluate the levels of cIAP1. Representative immunoblot of n = 3 mice per group. D) Percentage of superficial B16-F10 tumor nodules normalized to the average of the wildtype control. Tumor cells were injected *i.v*. and nodules formed in the lung were counted 13 days later (n = >3 experiments). E) Percentage of superficial B16-F10 tumor nodules normalized to the average of the wildtype recipient control. Tumor cells were injected *i.v*. and nodules formed in the lung were counted 12 days later (n = 3 experiments). Data are presented as mean ± SEM. *P < 0.05; **P < 0.01; ***P < 0.001, ns = not significant. 2-way ANOVA with Bonferroni’s test (A, B), 1-way ANOVA with Bonferroni’s test (D), twotailed unpaired t-test (E).

To confirm that loss of cIAP1 in the immune system does not interfere with tumor metastasis, we crossed *ciap1^fl/fl^ciap2^frt/frt^* with Vav-cre mice to deplete cIAP1 specifically in the hematopoietic system (*ciap1^vav^*). cIAP1 deletion was confirmed by protein detection in spleen and thymus (Figure 2C). B16-F10 melanoma cells were injected intravenously and the results showed that there was no reduction in the number of tumor colonies formed in the lungs of *ciap1^vav^* mice compared to *ciap1^fl/fl^* mice (Figure 2D).

To determine whether the loss of cIAP1 in the stromal compartment conferred reduction of lung tumor colonization, *ciap1^fl/fl^* and *ciap1^−/−^* mice were reconstituted with CD45.1 wildtype bone marrow. We observed a significant reduction of tumor nodules formed in the lungs of cIAP1 deficient recipients (Figure 2E). Together, the data suggest that cIAP1 is not essential for the recruitment of immune cells early upon tumor challenge and that the immune system deficient in cIAP1 at steady state does not influence the ability of tumor cells to home to the lung. Moreover, cIAP1 in the radio-resistant compartment promoted lung colonization.

### The *ciap1^−/−^* endothelium forms a stronger barrier against B16-F10 cell extravasation

To check whether the endothelium is the critical compartment *in vivo* for the reduction in the number of tumor colonies in the lungs of *ciap1^−/−^* mice, we crossed *ciap1^fl/fl^* with tamoxifen inducible VE-cadherin-cre::ER mice (*ciap1^VEC^*). Here, *ciap1^VEC^* mice showed reduced tumor colony numbers in the lung by approximately 50%, similar to complete *ciap1^−/−^* mice (Figure 3A), supporting the role of endothelium as the key player in our model. In the absence of cIAP1, cIAP2 up-regulation has been reported(29). However, combinatorial loss of cIAP1 and cIAP2 in the endothelium (*ciap1^VEC^ciap2^−/−^* mice) displayed a similar reduction in tumor load compared to *ciap1^VEC^ciap2^frt/frt^* mice, suggesting potential up-regulation of cIAP2 did not prevent tumor cell lung colonization. The efficiency of knocking out *ciap1* in the lung endothelium was confirmed by qPCR using primary isolated endothelial (CD31^+^) cells (75% reduction) (Figure 3B). Our previous data suggested that tumor growth is not hindered in *ciap1^−/−^* mice. Consequently, we focused on addressing whether tumor cells can transmigrate through the endothelial layer utilizing transmigration assays. The loss of cIAP1 in endothelial cells effectively reduced the number of transmigrating B16-F10 cells by 60% (Figure 3C). Apoptosis and necroptosis have been implicated in tumor transmigration(23, 30) and the loss of cIAP1 has been shown to sensitize cells to apoptosis. To determine whether tumor cells triggered the death of cIAP1 deficient endothelial cells, we co-cultured human umbilical vein endothelial cells (HUVECs) with various numbers of B16-F10 tumor cells and analyzed for cell death by flow cytometry. Using Birinapant in the media to simulate cIAP1 loss, we found no difference in the percentage of cell death in the HUVECs or the tumor cells, with or without Birinapant (Supplemental Figure 3A-3D). These results suggest a non-apoptotic function of cIAP1 in endothelial cells aids in tumor cell extravasation.

**Figure 3.**
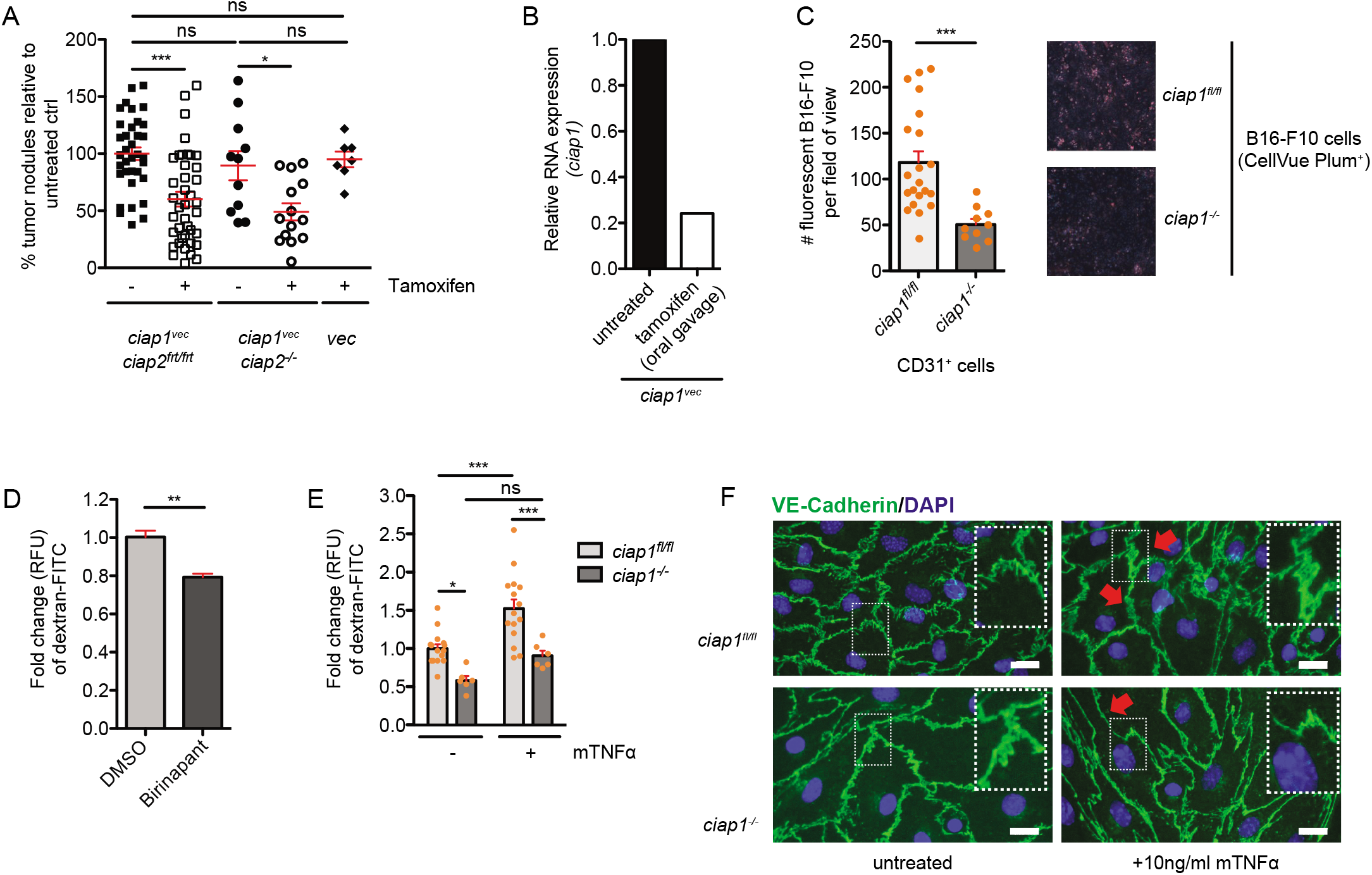
The *ciap1^−/−^* endothelium does not respond to permeability cues from B16-F10 tumor cells. A) Percentage of superficial B16-F10 tumor nodules normalized to the average of the untreated wildtype control. Tumor cells were injected *i.v*. into *ciap1^vec^* mice and controls. Nodules formed in the lung were counted 13 days later (n ≥ 3 experiments). B) Relative quantification of *ciap1* expression levels by qPCR. Total RNA was isolated from lung primary endothelial cells of untreated or tamoxifen-fed *ciap1^vec^* mice to induce recombination. C) Transendothelial migration of CellVue Plum-stained B16-F10 cells through a monolayer of primary isolated endothelial cells (n = 2 experiments, 2-4 biological replicates). Representative images of transwells demonstrating the amount of migrated CellVue Plum^+^ B16-F10 cells (red) surrounded by endothelial cells stained only with DAPI (blue). D, E) HUVEC monolayers were pre-treated with Birinapant for 16h (D) and primary isolated endothelial cell (EC) monolayers were stimulated with mouse TNF (mTNF) for 18h (E) before permeability to dextran-FITC was assessed (30min). Control cells were treated with DMSO or left untreated (n = 3 transwells/condition) (D); n = 2-5 biological replicates, 3 transwells/condition/experiment, 4 experiments (E)). F) Immunofluorescence staining of primary isolated ECs with DAPI (blue) and primary antibody against VE-Cadherin (green) treated with mTNF for 18h. Scale bars: 20 μm. Data are presented as mean ± SEM. *p < 0.05; **p < 0.01; ***p < 0.001, ns = not significant. 1-way ANOVA with Bonferroni’s test (A), two-tailed unpaired t-test (C, D), 2-way ANOVA with Bonferroni’s test (E).

Previous data have shown inhibition of XIAP and cIAP1/2 in HUVECs reduces permeability in response to thrombin(24). In agreement, we found the use of Birinapant reduced permeability of HUVECs *in vitro* (Figure 3D). Similarly, the permeability of primary *ciap1^−/−^* endothelial cell barrier was reduced compared to *ciap1^fl/fl^* cells without stimulation and the addition of mouse TNF (mTNF) resulted in an increase in permeability in *ciap1^fl/fl^* but not in *ciap1^−/−^* endothelial cells (Figure 3E). Endothelial cell purity after isolation was quantified and verified by endothelial cell specific markers, CD31^+^ and VE-cadherin staining (Supplemental Figure 4A, B). In response to mTNF, a known permeability factor, the junctions of VE-cadherin are disrupted in endothelial cells to promote permeability. In TNF treated *ciap1^−/−^* endothelial cells, VE-cadherin remained continuous and organized compared to *ciap1^fl/fl^* cells, where VE-cadherin staining became less distinct (Figure 3F). These results suggest cIAP1 regulates the morphology and response of endothelial cells to permeability factors and that the loss of cIAP1 specifically in adult endothelial cells leads to a decrease in permeability and extravasation of tumor cells.

### LTA but not TNF induces tumor cell extravasation via the TNFR1/cIAP1 axis

Since cIAP1 is well known to act downstream of the TNF/TNFRs pathway and TNF is a known cytokine involved in permeability, we examined whether TNF and TNFRs levels are influenced upon tumor challenge in the absence of cIAP1. Surprisingly, increasing levels of TNF could be detected in whole lung homogenates by luminex multiple assay 2h and 6h post tumor challenge, suggesting that TNF might be important for the extravasation of the tumor cells. While the levels of TNFR1 did not change, the levels of TNFR2 increased after tumor challenge (Figure 4A). Other angiogenic/permeability factors, such FGFb, did not show any fluctuation (Supplemental Figure 5).

**Figure 4.**
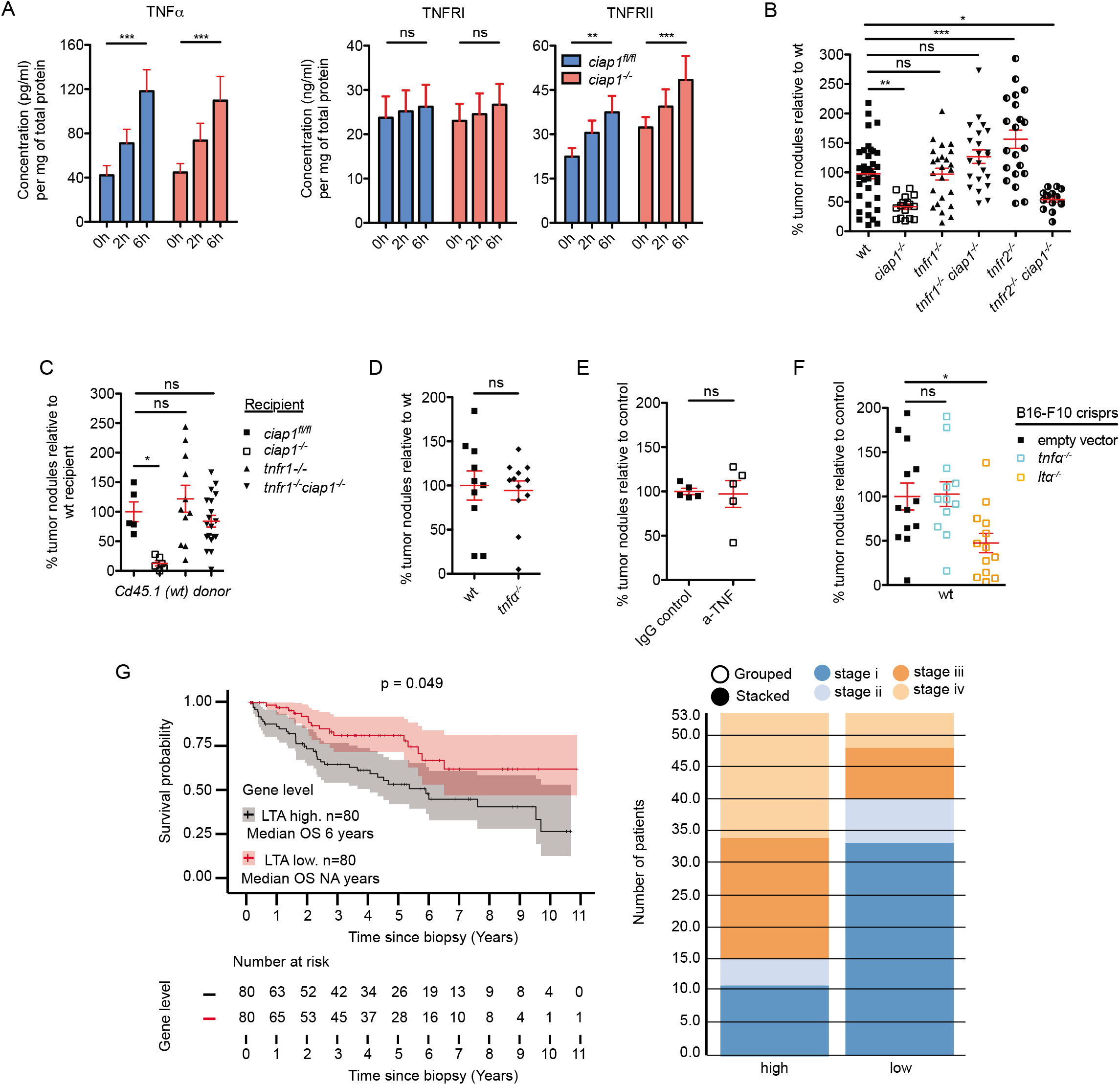
Lymphotoxin A (LTA) activates TNFR1 on the endothelium to induce tumor extravasation downstream of cIAP1. A) Levels of TNF and TNFRs assessed using multiplex analysis assay of total lung lysates (n = 2 experiments, 6 mice/group). B-E) Percentage of superficial B16-F10 tumor nodules normalized to the average of the respective controls. Tumor cells were injected *i.v*. into wildtype and transgenic mice (B, D), bone marrow chimeras (C) or antibody pre-treated wildtype mice (E), and nodules formed in the lung were counted 12 (C) or 13 days later (B, D-E). Data shown are pooled from n≥3 (B), 3 (C), 2 (D), 1 (E) experiments. F) Percentage of superficial B16-F10 tumor nodules normalized to the average of the respective control. LentiCRISPR-engineered B16-F10 cells against *tnf, lta* and control (empty vector) were injected *i.v*. into wildtype mice and nodules formed in the lung were counted 12 days later (n = 3 experiments). G) Survival curve of high (>85th percentile) versus low (<15th percentile) LTA mRNA expression in kidney renal clear cell carcinoma from TCGA data and distribution of high versus low expression of LTA by disease stage. Data are presented as mean ± SEM. *p < 0.05; **p < 0.01; ***p < 0.001, ns = not significant. 2-way ANOVA with Bonferroni’s test (A), 1-way ANOVA with Dunnett’s test (B, C, F), two-tailed unpaired t-test (D, E).

To assess the involvement of TNFRs upstream of cIAP1 in tumor extravasation we analyzed the number of tumor colonies formed in the lungs of *tnfr1^−/−^, tnfr2^−/−^, tnfr1^−/−^ciap1^−/−^* and *tnfr2^−/−^ciap1^−/−^* mice. Loss of TNFR1 did not affect tumor colonies in the lung, while the loss of TNFR2 displayed an increased number of tumor nodules. Interestingly, *tnfr1^−/−^ciap1^−/−^* had similar numbers of tumor nodules in the lung compared to wildtype mice, whereas *tnfr2^−/−^ ciap1^−/−^* mice mirrored the *ciap1^−/−^* phenotype (Figure 4B). Bone marrow chimera experiments further supported the combinatorial loss of TNFR1 and cIAP1 in the radio-resistant compartment was the cause for the restoration of the tumor load (Figure 4C). While TNFR1 has been identified as an important factor to induce endothelial permeability *in vitro* and *in vivo*, antibody blockade of TNFR2 hindered TNF-induced permeability in liver vessels, suggesting that TNFR2 can also contribute to the regulation of TNF-induced permeability *in vivo*(31). Together, these results suggest TNFR2 can compensate for the loss of TNFR1 even in the absence of cIAP1. *Tnfr2^−/−^* mice showed a higher number of tumor nodules in the lung, potentially because cIAP1 levels remain constant, supporting enhanced extravasation of tumor cells upon TNFR1 stimulation.

TNF and LTA are the known ligands for TNFR1 and TNFR2. We found the tumor load in *tnf^−/−^* mice was comparable to wildtype mice and injection of anti-TNF into the mice did not prevent tumor cells from extravasating and forming tumor nodules in the lung (Figure 4D and 4E). We then knocked out TNF or LTA in B16-F10 cells using CRISPR. Polyclonal populations selected by drug resistance were sequenced for genetic loss (Supplemental Figure 6A and 6B). Similar to our previous data, loss of TNF in B16-F10 cells did not affect tumor cell nodules in the lung but the loss of LTA caused a significant decrease in the tumor load (Figure 4F). These results suggest a mechanism in which B16-F10 tumor cell extravasation is mediated by the release of LTA and the subsequent activation of TNFRs.

To determine whether high expression of LTA and metastasis are associated, we correlated expression levels of LTA (mRNA) to cancer patient survival and disease stage using publicly available RNA expression data from The Cancer Genome Atlas (TCGA)(32). Kidney, lung cancer and melanoma are tumors known to metastasize to the lung. Among these tumor types, kidney renal clear cell carcinoma and papillary cell carcinoma displayed high expression of LTA mRNA (> 85^th^ quartile) that correlated with poor survival, while there was no correlation with TNF mRNA expression levels (Figure 4G, p<0.05 and Supplemental Figure 7A, B). We then segregated the high versus low expression by disease stage and found a great numbers of patients, categorized as disease stage 3 and 4, showing high expression of LTA. Surprisingly, in skin cutaneous melanoma and lung adenocarcinoma, the prognosis was the reverse, where high expression of LTA correlated with increased survival (Supplemental Figure 7C, D). Taken together, these results suggest a metastatic strategy which tumor cells utilize, involving release of LTA and activation of TNFRs. LTA as a biomarker for metastasis and survival in different types of cancer necessitates further study.

The complexity of understanding the metastatic cascade lies in the numerous interactions and cell types that are involved in this process. By focusing on the last stages of metastasis we were able to identify a novel role of Smac mimetics against tumor extravasation. This particularly pertains to tumor types that metastasize to the lung. In agreement, Smac mimetics (JP1584) in combination with TRAIL antagonism had been used in an orthotopic syngeneic rat model of hepatic cholangiocarcinoma leading to a reduction in extrahepatic metastasis(33). Therefore, the promise of Smac mimetics as therapeutic agents might lie not only on targeting primary tumors but also being used as metastasis-reducing compounds in certain tumor types.

## Materials and Methods

Animal work. Animals were maintained in specific pathogen free (SPF) and optimized hygiene conditions (OHB), and experiments were approved by Zürich Cantonal Veterinary Committee in accordance to the guidelines of the Swiss Animal Protection Law (License 119/2012 and 186/2015). *ciap1^fl/fl^ciap2^frt/frt^* (*ciap1^fl/fl^*), *ciap1^−/−^ciap2^frt/frt^* (*ciap1^−/−^*), *ciap1^fl/fl^ciap2^−/−^ (ciap2^−/−^*) mice were obtained from Dr. John Silke (WEHI, Melbourne, Australia); *Tnfr1^−/−^*, *tnfr2^−/−^* from Prof. Dr. Adriano Fontana and Adriano Aguzzi, respectively; Vav1-icre (vav) from Jackson Labs; Tg(Cdh5-cre/ERT2)1Rha (VEC) from Prof. Dr. Ralf Adams. Age and sex matched mice (6-12 weeks) were injected with B16-F10 (10^6^ cells/ml), MC-38 (1.5×10^6^ cells/ml), Lewis lung carcinoma (LLC) (1.5×10^6^ cells/ml) intravenously in 200 μl PBS or subcutaneous in 100 μl PBS. Mice were sacrificed by anesthesia, lungs perfused and blinded before further processing (nodule counting, flow cytometry analysis). Subcutaneous tumors were injected only in female mice (6-12 weeks old). For bone marrow chimeric experiments, mice were irradiated two times at 5.5Gy (11Gy total dose), 15h apart, injected with CD45.1 wildtype donor bone marrow, supplied with antibiotics for the first three weeks (neomycin into drinking water at 1mg/ml) and assessed for reconstitution efficiency at 6 weeks. Only mice with reconstitution efficiency >95% were used for further experiments.

For additional materials and methods, see Supplementary Materials & Methods

## Author Contributions

LV and WWW performed data analysis, designed the study concept and experiments and wrote the manuscript. KH and LMS performed experiments and aided in manuscript writing.

## Acknowledgements

We thank LASC animal technicians for animal husbandry. We also thank our research funding agencies for supporting this study: Peter Müller Fellowship (KH), Forschungskredit ‘CanDoc’ Universität Zürich Fellowship (KH, LV, LMS), Olga Mayer Stiftung (LMS), Sassella Stiftung and SNSF project grant (310030–138085, 310030-159613), Kurt and Senta-Hermann Stiftung, Krebsliga Schweiz (KFS-3386-02-2014), Zürich Krebsliga and Clöetta Medical Research fellow to WWLW. The authors acknowledge the assistance and support of the Center for Microscopy and Image Analysis, University of Zurich.

## Supplemental Figure Legends

**Supplemental Figure 1. Syngeniec tumor cells show varied responses to loss of cIAP1 *in vitro* and *in vivo***. A) Percentage of superficial B16-F10 tumor nodules normalized to the average of the captisol control. Wiltdype mice were injected i.p. with Birinapant or captisol before the tumor challenge. Tumor cells were injected i.v. and nodules formed in the lung were counted 13 days later (n = 2 experiments). The experimental set-up is illustrated below the graphs. B) B16-F10 melanoma, MC-38 GFP colon carcinoma and Lewis lung carcinoma (LLC) cells were treated with Smac mimetics (Birinapant or 12911), mouse TNF and the caspase inhibitor QVD-OPh. DMSO was used as a control. Cell death was assessed by propidium iodide (B16-F10, LLC) or fixable viability dye eFluor780 uptake using flow cytometry (n = 3-4, 3 experiments). C) Percentage of superficial LLC tumor nodules normalized to the average of the wt control. Tumor cells were injected i.v. and nodules formed in the lung were counted 21 days later (n = 2 experiments). Data are presented as mean ± SEM. *p < 0.05; **p < 0.01; ***p < 0.001, ns = not significant. Two-tailed unpaired t-test (A), 2-way ANOVA with Bonferroni’s test (B), 1-way ANOVA with Bonferroni’s test (C).

**Supplemental Figure 2. Lung myeloid and lymphoid immune infiltrate kinetics is similar between wildtype and *ciap1^−/−^* mice subjected to tumor challenge with B16-F10 cells.** Analysis of lung immune cell infiltrates following B16-F10 tumor challenge by flow cytometry. Interstitial macrophages (MΦ) (CD11b^high^MHCII^+^CD11c^+^CD64^high^), eosinophils (CD11b^high^CD11c^−^SiglecF^+^), CD4^+^ T cells (CD3^+^CD4^+^) and CD8^+^ T cells (CD3^+^CD8^+^) were pre-gated on singlets, live and CD45^+^ cells (n = 2 experiments, 6 mice/group). B-D) Levels of matrix metalloproteinase-related factors (B), cell adhesion molecules (C) and cytokines/chemokines (D) assessed using multiplex analysis assay of total lung lysates (n = 2 experiments, 6 mice/group). Data are presented as mean ± SEM. *p < 0.05; **p < 0.01; ***p < 0.001. 2-way ANOVA with Bonferroni’s test.

**Supplemental Figure 3. Tumor cells and Smac mimetics do not induce the death of HUVEC cells in vitro**. A-D) HUVEC monolayers were co-cultured with increasing numbers of CellVue Plum-stained B16-F10 or MC-38 GFP cells, with or without Birinapant, for 24h. Cell death of HUVECs (A, C) and tumor cells (B, D) was assessed by propidium iodide (PI) (A, B) or fixable viability eFluor780 dye (C, D) uptake using flow cytometry (n = 3-4, 3 experiments (A, B)). Data are presented as mean ± SEM. *p < 0.05; **p < 0.01; ***p < 0.001. 1-way ANOVA with Bonferroni’s test (A, B).

**Supplemental Figure 4. Purity of isolated primary pulmonary endothelial cells (ECs) following magnetic-activated cell sorting (MACS)**. A) Primary isolated pulmonary ECs (CD31+) were analysed by flow cytometry following MACS separation. Cells from the elution, enriched in CD31+ cells, and flowthrough were stained with an a-rat-FITC secondary antibody and propidium iodide (PI) to assay for live cells. Pre-gated on singlets. Representative FACS plots. B) Immunofluorescence staining of primary isolated ECs with DAPI (blue) and primary antibody against VE-Cadherin (green). Representative picture. Scale bar: 100 μm.

**Supplemental Figure 5. Growth factor kinetics do not show a profound difference between wildtype and *ciap1^−/−^* mice subjected to tumor challenge with B16-F10 cells**. Levels of growth factors assessed using multiplex analysis assay of total lung lysates (n = 2 experiments, 6 mice/group). Data are presented as mean ± SEM. *p < 0.05; **p < 0.01; ***p < 0.001. 2-way ANOVA with Bonferroni’s test.

**Supplemental Figure 6. Design of B16-F10 lenti-crisprs and sequence analysis**. A,B) Guide RNA (gRNA) primers for *tnf* (A) and *lta* (B) were designed to bind in proximity and downstream of the start codon. Representative sequencing results of the polyclonal lenticrispr populations and signle clones are shown aligned to the coding strands of the genes. Mutagenesis is apparent taking place around 4-5 nucleotides upstream of the protospacer adjacent motif (PAM) sequence shown in red.

**Supplemental Figure 7. Correlation between survival of cancer patients and expression levels of LTA and TNF**. A-D) Survival curves of high (>85th percentile) versus low (<15th percentile) LTA and TNF mRNA expression in kidney renal clear cell carcinoma (KIRC), kidney renal papillary cell carcinoma (KIRP), skin cutaneous melanoma (SKCM) and lung adenocarcinoma (LUAD) from TCGA data.

## References

1. Zender L et al. Identification and validation of oncogenes in liver cancer using an integrative oncogenomic approach. Cell 2006;125(7):1253–1267.

2. Dai Z et al. A comprehensive search for DNA amplification in lung cancer identifies inhibitors of apoptosis cIAP1 and cIAP2 as candidate oncogenes. Hum Mol Genet 2003;12(7):791–801.

3. Imoto I et al. Identification of cIAP1 as a candidate target gene within an amplicon at 11q22 in esophageal squamous cell carcinomas. Cancer Res 2001;61(18):6629–6634.

4. Imoto I et al. Expression of cIAP1, a target for 11q22 amplification, correlates with resistance of cervical cancers to radiotherapy. Cancer Res 2002;62(17):4860–4866.

5. Du CY, Fang M, Li YC, Li L, Wang XD. Smac, a mitochondrial protein that promotes cytochrome c-dependent caspase activation by eliminating IAP inhibition. Cell 2000;102(1):33–42.

6. Verhagen AM et al. Identification of DIABLO, a mammalian protein that promotes apoptosis by binding to and antagonizing IAP proteins. Cell 2000;102(1):43–53.

7. Wu G et al. Structural basis of IAP recognition by Smac/DIABLO. Nature 2000;408(6815):1008–1012.

8. Vince JE et al. IAP antagonists target cIAP1 to induce TNFalpha-dependent apoptosis. Cell 2007;131(4):682–693.

9. Varfolomeev E et al. IAP antagonists induce autoubiquitination of c-IAPs, NF-kappaB activation, and TNFalpha-dependent apoptosis. Cell 2007;131(4):669–681.

10. Gaither A et al. A Smac mimetic rescue screen reveals roles for inhibitor of apoptosis proteins in tumor necrosis factor-alpha signaling. Cancer Res 2007;67(24):11493–11498.

11. Petersen SL et al. Autocrine TNFalpha signaling renders human cancer cells susceptible to Smac-mimetic-induced apoptosis. Cancer Cell 2007;12(5):445–456.

12. Siegmund D, Kums J, Ehrenschwender M, Wajant H. Activation of TNFR2 sensitizes macrophages for TNFR1-mediated necroptosis. Cell Death Dis 2016;7(9):e2375–e2375.

13. Knop J et al. TNFR2 induced priming of NLRP3-inflammasome via RIPK1 leads to pyroptosis in XIAP deficient cells. bioRxiv 2019;:550749.

14. Lecis D et al. Smac mimetics induce inflammation and necrotic tumour cell death by modulating macrophage activity. Cell Death Dis 2013;4(11):e920.

15. Beug ST et al. Smac mimetics and innate immune stimuli synergize to promote tumor death. Nature Biotechnology 2014;32(2):182–190.

16. Chesi M et al. IAP antagonists induce anti-tumor immunity in multiple myeloma. Nature Medicine [published online ahead of print: November 14, 2016]; doi:10.1038/nm.4229

17. Kearney CJ et al. PD-L1 and lAPs co-operate to protect tumors from cytotoxic lymphocyte-derived TNF 2017;24(10):1705–1716.

18. Beug ST et al. Smac mimetics synergize with immune checkpoint inhibitors to promote tumour immunity against glioblastoma. Nature Communications 2017;8:1–15.

19. Witt A et al. IAP antagonization promotes inflammatory destruction of vascular endothelium. EMBO Rep 2015;16(6):719–727.

20. Strilic B, Offermanns S. Intravascular Survival and Extravasation of Tumor Cells. Cancer Cell 2017;32(3):282–293.

21. Santoro MM, Samuel T, Mitchell T, Reed JC, Stainier DYR. Birc2 (clap1) regulates endothelial cell integrity and blood vessel homeostasis. Nat Genet 2007;39(11):1397–1402.

22. Moulin M et al. lAPs limit activation of RIP kinases by TNF receptor 1 during development. EMBO J 2012;31(7):1679–1691.

23. Strilic B et al. Tumour-cell-induced endothelial cell necroptosis via death receptor 6 promotes metastasis. Nature 2016;536(7615):215–218.

24. Hornburger MC et al. A novel role for inhibitor of apoptosis (IAP) proteins as regulators of endothelial barrier function by mediating RhoA activation. The FASEB Journal 2014;28(4):1938–1946.

25. Hänggi K et al. RIPK1/RIPK3 promotes vascular permeability to allow tumor cell extravasation independent of its necroptotic function. [Internet]. Cell Death Dis 2017;8(2):e2588.

26. Qian B-Z et al. CCL2 recruits inflammatory monocytes to facilitate breast-tumour metastasis. Nature 2011;475(7355):222–225.

27. Wolf MJ et al. Endothelial CCR2 signaling induced by colon carcinoma cells enables extravasation via the JAK2-Stat5 and p38MAPK pathway. Cancer Cell 2012;22(1):91–105.

28. Houghton AMG et al. Macrophage elastase (matrix metalloproteinase-12) suppresses growth of lung metastases [Internet]. Cancer Res 2006;66(12):6149–6155.

29. Conze DB et al. Posttranscriptional downregulation of c-IAP2 by the ubiquitin protein ligase c-IAP1 in vivo. Molecular and Cellular Biology 2005;25(8):3348–3356.

30. Kebers F et al. Induction of endothelial cell apoptosis by solid tumor cells. Experimental Cell Research 1998;240(2):197–205.

31. Ferrero E et al. Roles of tumor necrosis factor p55 and p75 receptors in TNF-alpha-induced vascular permeability. Am. J. Physiol., Cell Physiol. 2001;281(4):C1173–C1179.

32. Cheng PF, Dummer R, Levesque MP. Data mining The Cancer Genome Atlas in the era of precision cancer medicine. Swiss Med Wkly 2015;145:w14183.

33. Fingas CD et al. A smac mimetic reduces TNF related apoptosis inducing ligand (TRAIL)-induced invasion and metastasis of cholangiocarcinoma cells. Hepatology 2010;52(2):550–561.

