## Supplemental materials and methods, Supplemental Figures for "Tumor cell-derived lymphotoxin alpha triggers metastatic extravasation through TNFRs/cIAP1"

**Supplementary Materials & Methods**

**Cell culture**

Tumor cells were cultured at maximum 80% confluency in high glucose (4.5 g/l) Dulbecco's modified Eagle's medium (DMEM) (MC-38 GFP, LLC) or RPMI 1640 (Gibco, ThemoFisher Scientific, Germany) (B16-F10) supplemented with 10% fetal bovine serum (FBS; Gibco, ThemoFisher Scientific) and penicillin/streptomycin/glutamine (PSG, Gibco, ThemoFisher Scientific). For B16-F10 cells, media was switched to high glucose DMEM 48 h prior to injection to stimulate melanin production. Primary lung ECs and HUVECs (C-12200, Promocell, Germany) were cultured on gelatin pre-treated plates (0.1% gelatin) in EBM-2 media supplemented with the EGM-2 SingleQuot Kit according to the manufacturers recommendation (Lonza, Germany) or the equivalent Endopan 3 medium (PAN Biotech, Germany).

**Primary lung EC isolation**

Lungs were perfused with PBS and HBSS+Collagenase IV (1 mg/ml, Sigma Aldrich, Switzerland) and minced before incubation with HBSS+Collagenase IV for 45 min at 37 $^{o}$C and a cell suspension was obtained. Mouse lung cell suspension was incubated with anti-CD31 antibody (BioLegend, clone 390) for 30 min in PBS+0.5% BSA at 1:25 dilution in order to isolate lung ECs, followed by incubation with anti-rat IgG magnetic particles and sorted on LS columns according to the manufacturers recommendations (Miltenyi, Switzerland).

**Transendothelial migration assay**

Only EC populations of >90% purity (as assessed by CD31$^{+}$) were used. Briefly, 3x10$^{4}$ primary ECs were seeded 3-4 days after isolation on a gelatin coated insert (Corning, Netherlands, 8 µm pore size) in EBM-2 (Lonza) or Endopan3 (Pan Biotech). Two days after seeding, cell culture media was changed to RPMI 1640 (Gibco) in the bottom well (3% FBS) and insert (1% FBS) and 2x10$^{4}$ CellVue Plum$^{+}$ (Polysciences, Germany) membrane stained B16-F10 tumor cells were added in the insert. After 20h, the remaining cells inside the insert were removed, the membrane was fixed in 2% PFA and mounted on glass slides using SlowFade Gold with DAPI (Thermoscience Fisher, Germany). Microscopic fluorescence picture were taken at 10x magnification using ZEISS AXIO Scan Z.1 at the Centre for Microscopy and Image Analysis (ZMB), Switzerland. CellVue Plum$^{+}$ cells were counted per field of view (pictures was divided into 6x6 tiles and only the ten central tiles representing the insert were considered for counting).

**Dextran-FITC permeability assay**

3x10$^{4}$ HUVEC or primary ECs were seeded 3–4 days after isolation on a gelatin-coated insert (Corning, 8 µm pore size) in EBM-2 (Lonza) or Endopan3 (PAN Biotech) and grown to a confluent monolayer for 3 days. The medium was gradually replaced to ECGS medium (low glucose DMEM + 10% FBS + PSG + ascorbic acid (EBM-2 supplement, Lonza) + heparin (EBM-2 supplement, Lonza) + 30µg/ml endothelial growth supplement from bovine pituitary (ECGS, Sigma)). Cells were then pre-treated with mouse TNF or Birinapant for 18 h before dextran-FITC (70kDa, Sigma) was added into the insert at final concentration of 1 mg/ml. Supernatant from the bottom well was collected 30min after addition of dextran-FITC and fluorescence intensity was measured by TECAN Infinite 200 Pro multireader (Switzerland) using λEx = 490 nm and λEm = 520 nm.

**Multiplex assay**

Total lung lysates were prepared according to the manufacturer's instructions. A customized mouse Magnetic Luminex assay kit was used (R&D Systems). The measurements were performed on a Bio-Rad BioPlex 200 system (Bio-Rad Laboratories, Switzerland) and data were analyzed with the Bio-Plex software 6.0.

**Antibodies and viability dyes**

The following antibodies/viability dyes were used: CD11b-PE/Cy7 (M1/70), CD45-BV605 (30F-11), Ly6C-BV711 (HK1.4), CD11c-APC/Cy7 (N418), Ly6G-PerCP/Cy5.5 (1A8), MHCII-AF700 (M5/114.15.2), CD64-APC (X54-5/7.1), CD3-BV785 (17A2), CD4-APC (GK1.5), CD8-PerCP (YTS156.7.7), NK1.1-BV650 (PK136) from Biolegend; CD45-AF700 (30F-11), CD24-PE (M1/69), CD103-FITC (2E7), fixable viability dye eFluor506 and eFluor780 from eBioscience; CD144 (VE-cadherin, 11D4.1), SiglecF-BV421 (E50-2440), CD11b-BV605 (M1/70) from BD Biosciences; donkey anti-rat IgG (H+L)-AF488 from Jackson; propidium iodide (PI), beta-actin (AC-15) from Sigma; anti-cIAP1 (1E1-1-10) from ENZO Life Sciences, Germany and donkey anti-mouse and anti-rabbit IgG (H+L)-HRP from SouthernBiotech.

**Ligands and inhibitors**

Various ligands and inhibitors were used at the following concentrations: Mouse and human TNF (10ng/ml, purified from mammalian expression system pcDNA5 FRT fc hs TNFa and pCR3 fc ps mm TNF), Birinapant (500nM, Chemietek), Compound A (12911, 500nM, Tetralogics), QVD-OPh (5µM, Adipogen).

**Compounds used *in vivo***

Mice received an intraperitoneal injection of 200µg of IgG control or anti-TNF antibodies (MP6-XT22, Biolegend) in 100µl PBS 2.5h before the tumor challenge ($10^5$ B16-F10 cells in 100µl of PBS). Birinapant was dissolved in 12.5% captisol (5mg/ml, Capitsol, USA) and used for intraperitoneal injections at 5mg/Kg after 1:10 dilution in PBS. Tamoxifen (MedChemExpress, Sweden) was dissolved in corn oil (Sigma) at 20mg/ml. Young mice (5 weeks old) were fed for 3 consecutive days by oral gavage (150mg/kg).

**Western blotting**

Organs (lung, spleen, thymus) were lysed using DISC lysis buffer (20mM Tris-HCl pH 7.5, 150mM NaCl, 10% (v/v) glycerol, 1% (v/v) NP-40, 2mM EDTA, 5mM EGTA, 30mM NaF, 40mM b-glycerophosphate pH 7.2, 10mM sodium pyrophosphate, 2mM activated sodium orthovanadate, protease inhibitors). The insoluble fraction of the lysate was pelleted by centrifugation and removed. Lysates were boiled and run on 4-20% Bis-Tris Gel NuPAGE using MOPS buffer (Invitrogen). Proteins were then transferred onto PVDF-membrane (0.2µm, Thermo Scientific) using the Trans-Blot® Turbo™ Transfer System (Bio Rad), according to the manufacturer's instruction. After blocking with PBST containing 5% (w/v) skim milk, membranes were incubated with the indicated primary antibody in PBST containing 5% skim milk overnight at 4°C, followed by incubation with the respective secondary antibody. Protein level expression was visualized on a film (Kodak) using chemiluminescence (WesternBright ECL, Advansta).

**qPCR**

RNA from primary isolated lung ECs was extracted using Genezol (Geneaid, Taiwan) according to the manufacturer's instructions. cDNA was produced using MultiScribe™ Reverse Transcriptase and SYBR Green qPCR master mix (Thermo Fisher Scientific) was used for running the qPCR. Melting curves showed that single products were formed. The following primers were used: mciap1 (5': AAT GGT TTC CAA GGT GTG AG; 3': GGA CAA CAG CTG CTC AAG); B2M (5': TGG TGC TTG TCT CAC TGA CC; 3' CCG TTC TTC AGC ATT TGG AT). Relative standard curve analysis was performed using the housekeeping gene B2M and ECs from non-tamoxifen fed mouse were used as a calibrator for fold-change.

**Lentiviral CRISPR/Cas9 constructs**

The LentiCRISPRv2 one vector system was utilized to target the *tnf* and *lta* genomic loci in B16-F10 melanoma cells. The following oligos were used in order to clone the desired target sequence into the vector: mlta crispr forward (5'-CACCGGGGTGCCAAGCACCCTCAAG-3'), mlta reverse (5'-AAACCTTGAGGGTGCTTGGCACCCC-3'), mtnf forward (5'-CACCGAGAAAGCATGATCCGCGACG-3') and mtnf reverse (5'-AAACCGTCGCGGATCATGCTTTCTC-3'). Oligos were designed using the Benchling CRISPR gRNA Design tool (http://www.benchling.com). For cloning, viral production, tumor cell viral transduction and selection, we followed the recommended guidelines(1, 2).

### Statistical analysis

All data is presented in mean ± SEM. Figures were prepared in Illustrator CC 2015 (Adobe) and Prism 5 (GraphPad Software). Significance between genotypes and treatments was assessed by Student-t test, one-way or two-way ANOVA with * $p<0.05$, **$p<0.01$, ***$p<0.001$, ns = not significant using Prism 5.

### References


1. Sanjana NE, Shalem O, Zhang F. Improved vectors and genome-wide libraries for CRISPR screening. *Nature Methods* 2014;11(8):783–784.

2. Shalem O et al. Genome-scale CRISPR-Cas9 knockout screening in human cells. *Science* 2014;343(6166):84–87.

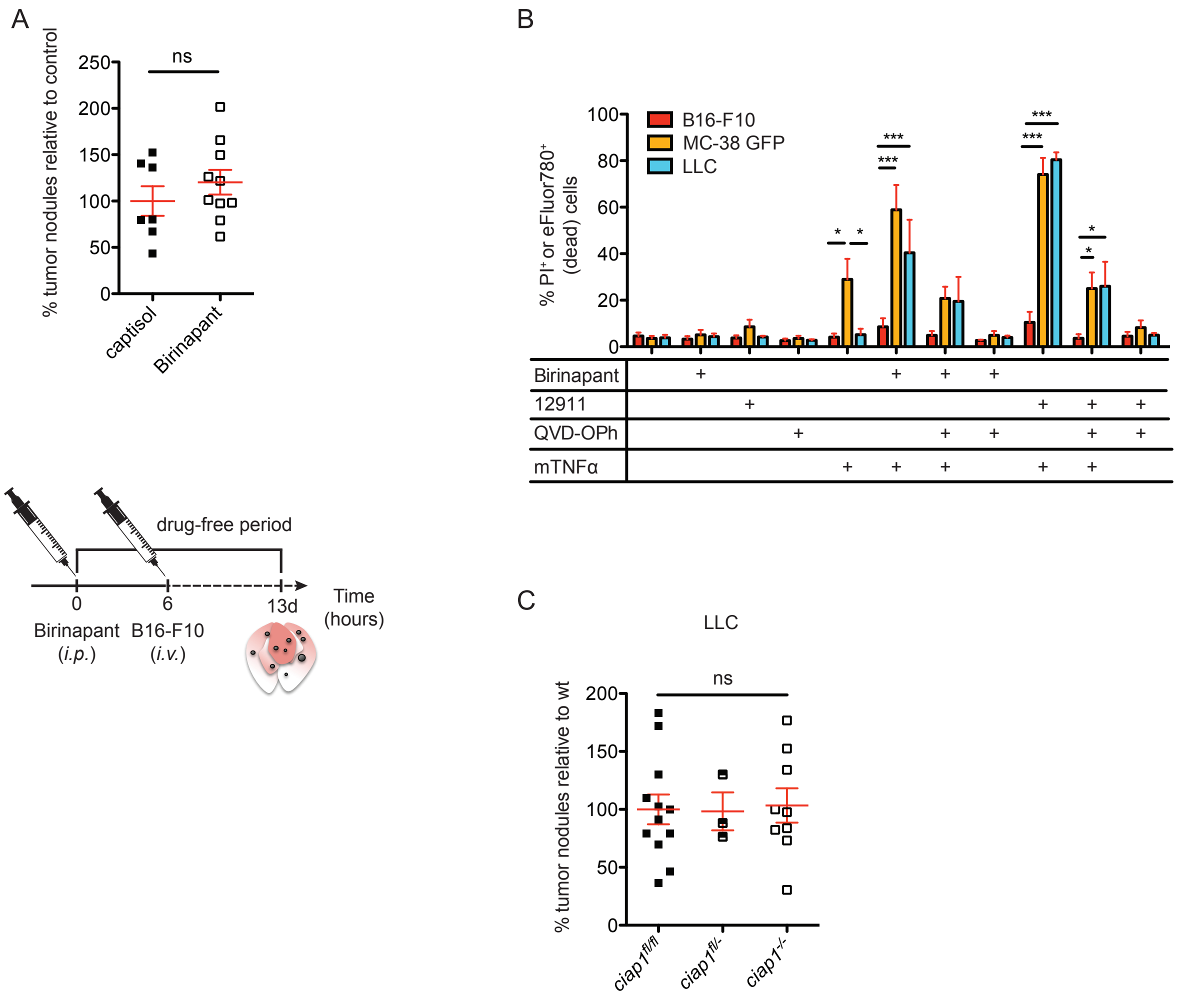


**Supplemental Figure 1. Syngeniec tumor cells show varied responses to loss of cIAP1 *in vitro* and *in vivo*.**

A) Percentage of superficial B16-F10 tumor nodules normalized to the average of the captisol control. Wiltdype mice were injected *i.p.* with Birinapant or captisol before the tumor challenge.Tumor cells were injected *i.v.* and nodules formed in the lung were counted 13 days later (n = 2 experiments). The experimental set-up is illustrated below the graphs. B) B16-F10 melanoma, MC-38 GFP colon carcinoma and Lewis lung carcinoma (LLC) cells were treated with Smac mimetics (Birinapant or 12911), mouse TNF and the caspase inhibitor QVD-OPh. DMSO was used as a control. Cell death was assessed by propidium iodide (B16-F10, LLC) or fixable viability dye eFluor780 uptake using flow cytometry (n = 3-4, 3 experiments). C) Percentage of superficial LLC tumor nodules normalized to the average of the wt control. Tumor cells were injected *i.v.* and nodules formed in the lung were counted 21 days later (n = 2 experiments). Data are presented as mean ± SEM. *p < 0.05; **p < 0.01; ***p < 0.001, ns = not significant. Two-tailed unpaired t-test (A), 2-way ANOVA with Bonferroni's test (B), 1-way ANOVA with Bonferroni's test (C).

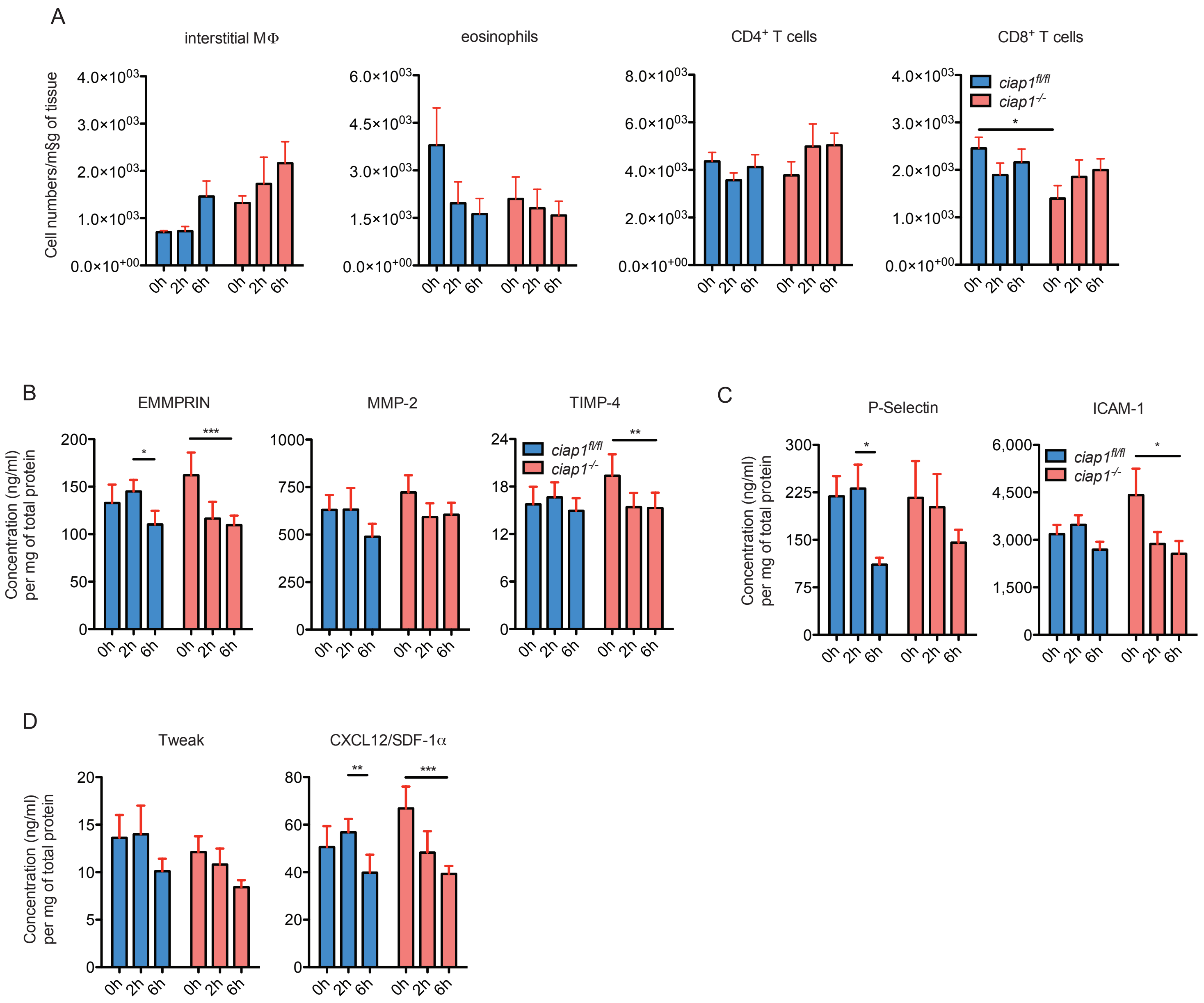


**Supplemental Figure 2. Lung myeloid and lymphoid immune infiltrate kinetics is similar between wildtype and *ciap1*$^{-/-}$ mice subjected to tumor challenge with B16-F10 cells.**

A) Analysis of lung immune cell infiltrates following B16-F10 tumor challenge by flow cytometry. Interstitial macrophages (MΦ) (CD11b$^{high}$ MHCII$^{+}$CD11c$^{+}$CD64$^{high}$), eosinophils (CD11b$^{high}$CD11c$^{-}$SiglecF$^{+}$), CD4$^{+}$ T cells (CD3$^{+}$CD4$^{+}$) and CD8$^{+}$ T cells (CD3$^{+}$CD8$^{+}$) were pre-gated on singlets, live and CD45$^{+}$ cells (n = 2 experiments, 6 mice/group). B-D) Levels of matrix metalloproteinase-related factors (B), cell adhesion molecules (C) and cytokines/chemokines (D) assessed using multiplex analysis assay of total lung lysates (n = 2 experiments, 6 mice/group). Data are presented as mean ± SEM. *$p < 0.05$; **$p < 0.01$; ***$p < 0.001$. 2-way ANOVA with Bonferroni's test.

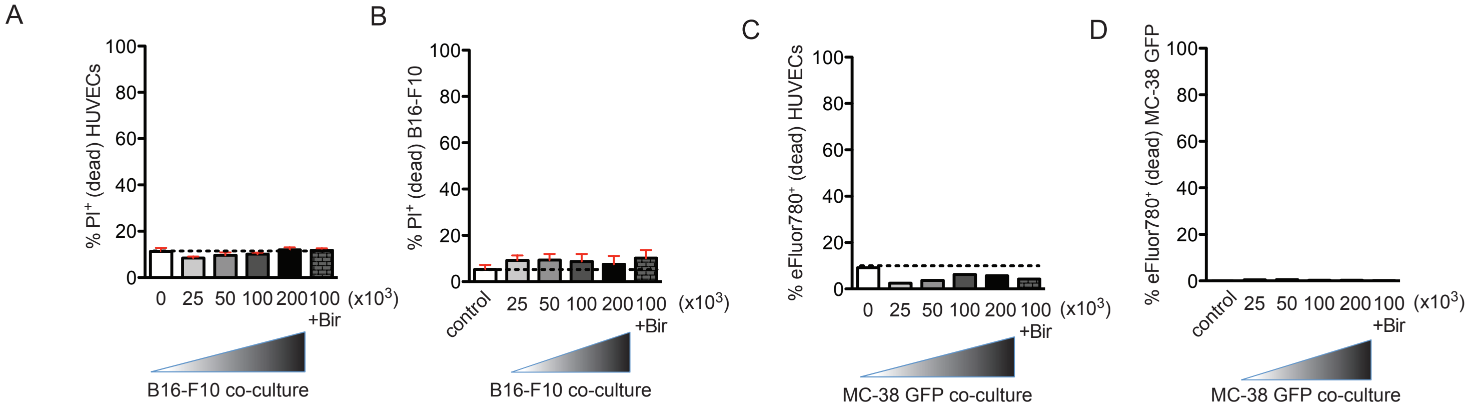


**Supplemental Figure 3. Tumor cells and Smac mimetics do not induce the death of HUVEC cells *in vitro*.**

A-D) HUVEC monolayers were co-cultured with increasing numbers of CellVue Plum-stained B16-F10 or MC-38 GFP cells, with or without Birinapant, for 24h. Cell death of HUVECs (A, C) and tumor cells (B, D) was assessed by propidium iodide (PI) (A, B) or fixable viability eFluor780 dye (C, D) uptake using flow cytometry (n = 3-4, 3 experiments (A, B)). Data are presented as mean ± SEM. *p < 0.05; **p < 0.01; ***p < 0.001. 1-way ANOVA with Bonferroni's test (A, B).

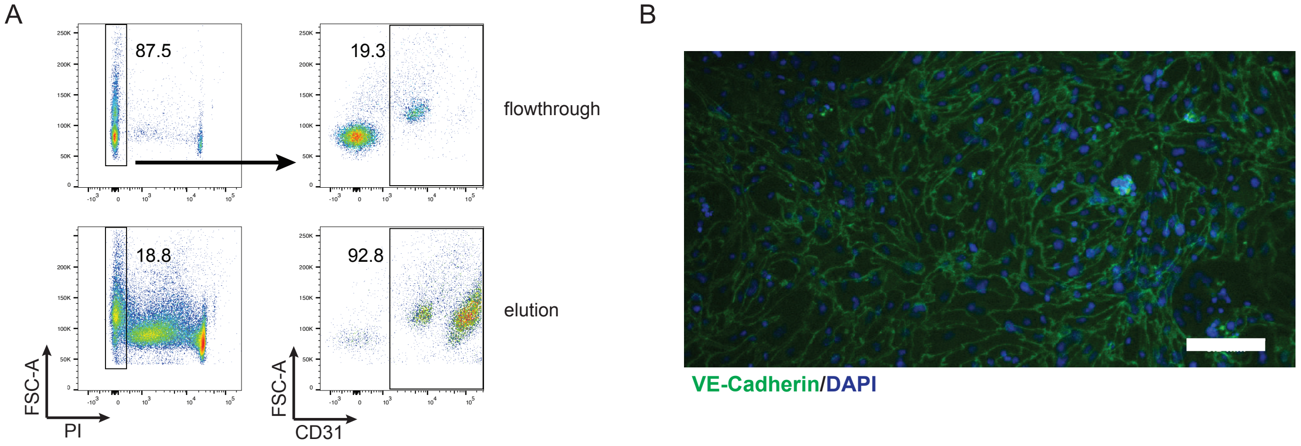


**Supplemental Figure 4. Purity of isolated primary pulmonary endothelial cells (ECs) following magnetic-activated cell sorting (MACS).**

A) Primary isolated pulmonary ECs ($CD31^+$) were analysed by flow cytometry following MACS separation. Cells from the elution, enriched in $CD31^+$ cells, and flowthrough were stained with an a-rat-FITC secondary antibody and propidium iodide (PI) to assay for live cells. Pre-gated on singlets. Representative FACS plots. B) Immunofluorescence staining of primary isolated ECs with DAPI (blue) and primary antibody against VE-Cadherin (green). Representative picture. Scale bar: 100 µm.

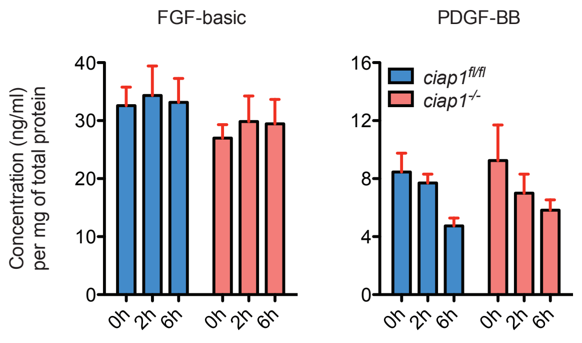


**Supplemental Figure 5. Growth factor kinetics do not show a profound difference between wt and *ciap1$^{-/-}$* mice subjected to tumor challenge with B16-F10 cells.**

Levels of growth factors assessed using multiplex analysis assay of total lung lysates (n = 2 experiments, 6 mice/group). Data are presented as mean ± SEM. *p < 0.05; **p < 0.01; ***p < 0.001. 2-way ANOVA with Bonferroni's test.

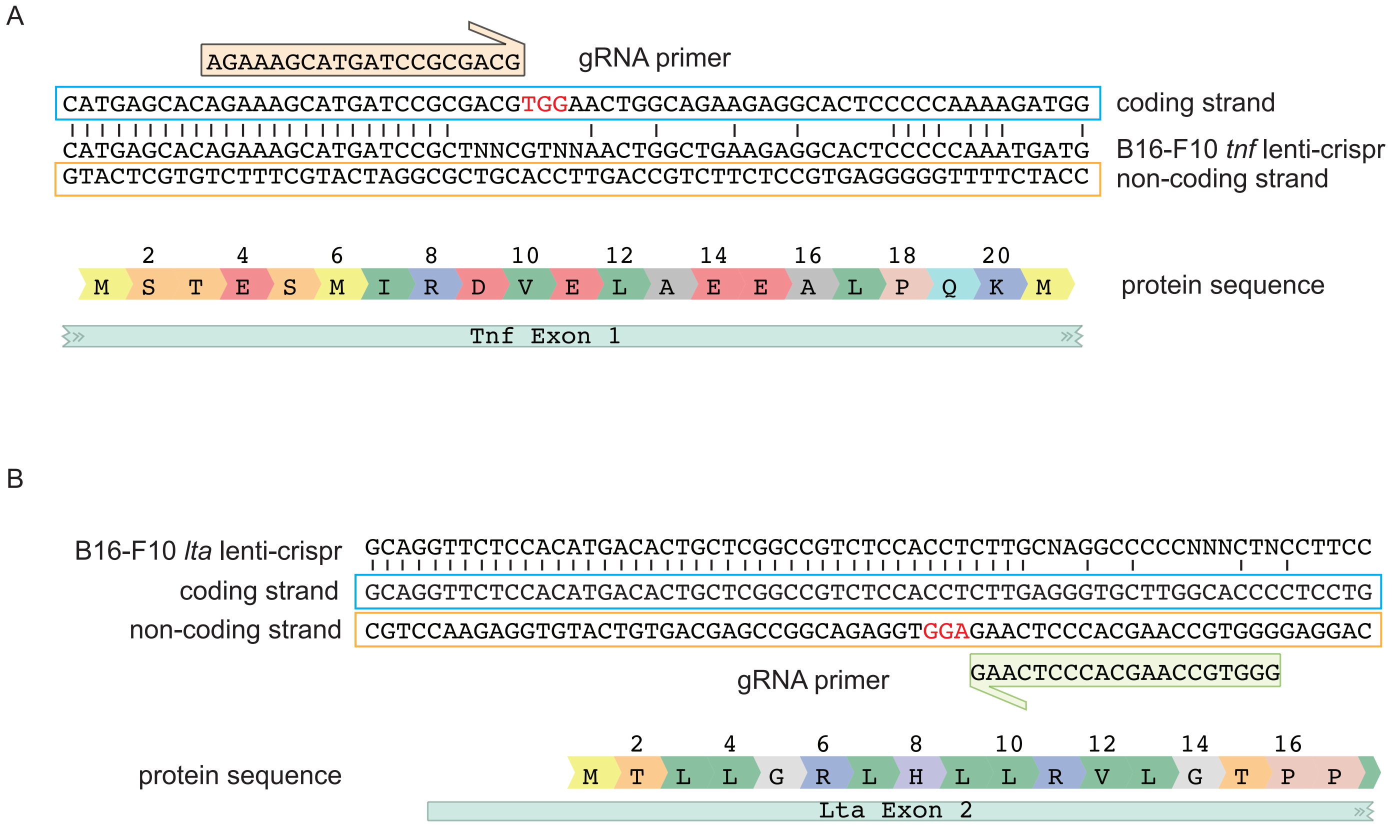


**Supplemental Figure 6. Design of B16-F10 lenti-crisprs and sequence analysis.**

A,B) Guide RNA (gRNA) primers for *tnf* (A) and *lta* (B) were designed to bind in proximity and downstream of the start codon. Representative sequencing results of the polyclonal lenti-crispr populations and signle clones are shown aligned to the coding strands of the genes. Mutagenesis is apparent taking place around 4-5 nucleotides upstream of the protospacer adjacent motif (PAM) sequence shown in red.

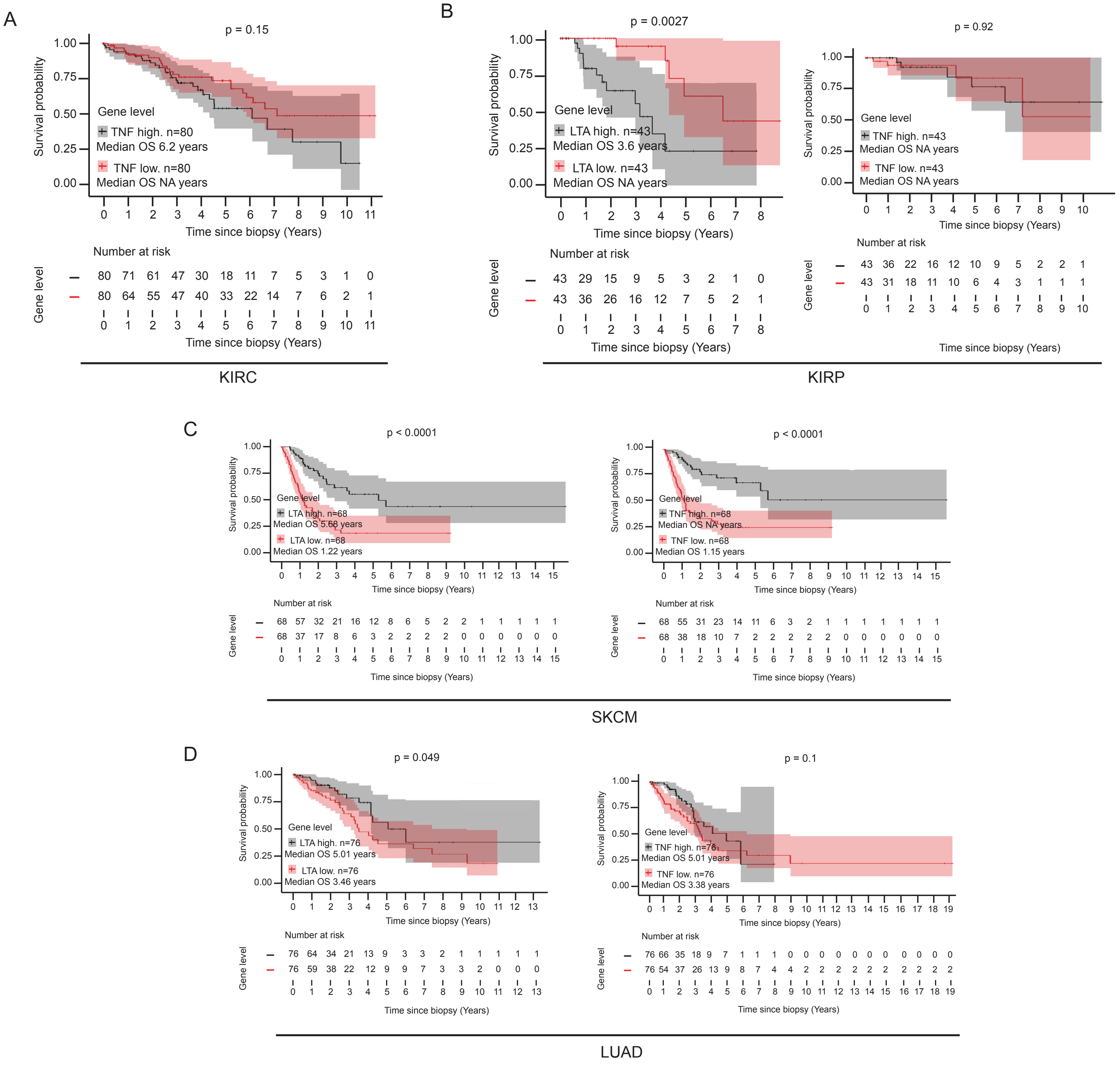


**Supplemental Figure 7. Correlation between survival of cancer patients and expression levels of LTA and TNF.**

A-D) Survival curves of high (>85th percentile) versus low (<15th percentile) LTA and TNF mRNA expression in kidney renal clear cell carcinoma (KIRC), kidney renal papillary cell carcinoma (KIRP), skin cutaneous melanoma (SKCM) and lung adenocarcinoma (LUAD) from TCGA data.

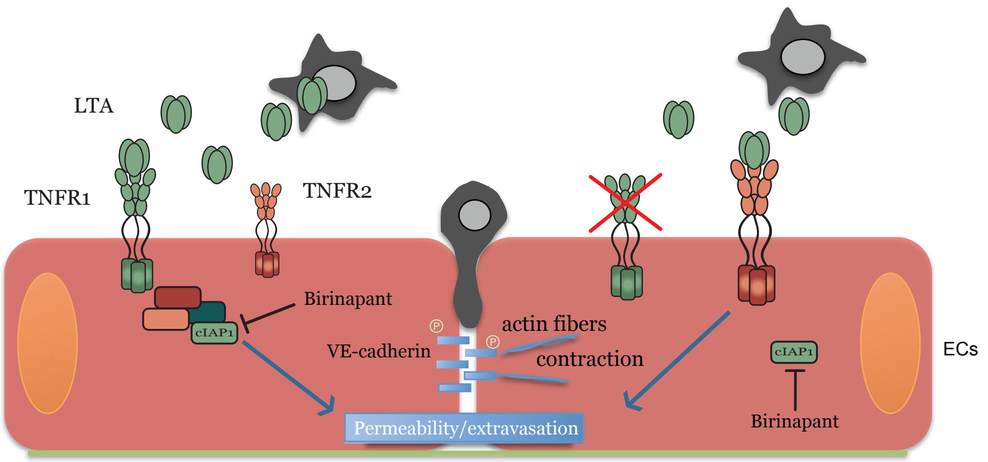


**Supplemental Figure 8. Graphical abstract.**

B16-F10 tumor cells secrete LTA, which activates primarily TNFR1 to induce extravasation. cIAP1 acts downstream of TNFR1 to modulate NF-κB activation and might also orchestrate the phosphorylation of VE-cadherin. When TNFR1 is absent or blocked, TNFR2 activation can induce the permeability without the need of cIAP1. Birinapant treatment can block tumor cell extravasation by inbiting cIAP1.
